# Mental maps without vision: Neural signatures of cognitive maps based on haptic input in the hippocampal formation

**DOI:** 10.1101/2023.10.20.563338

**Authors:** Loes Ottink, Lennard van den Berg, Imke Peters, Thea van der Geest, Koen Haak, Christian Doeller, Richard van Wezel

## Abstract

The human hippocampus is the key region for forming cognitive maps of our environment. Such a map can support spatial navigation. It is unclear whether this area is similarly involved when an environment is explored with our haptic sense. In this study, we investigated the neural representation of distances on a tactile map in the hippocampal formation, in visually impaired and sighted persons. To this end, 47 participants (22 persons with a visual impairment, PVIs, and 25 sighted controls) performed a navigation task where they learned a tactile city-like map including five item locations. We combined magnetic resonance imaging with adaptation analysis to assess representation of distances between item locations in the hippocampus and entorhinal cortex. Additionally, we assessed cognitive map formation on a behavioural level. We also looked at functional connectivity between navigation-related areas during a subsequent resting-state block. Our data reveal across all participants that the left entorhinal cortex represents distances between locations on a tactile map. Here, we provide the first evidence that maps in the hippocampal formation is preserved when an environment is presented in a non-visual modality. The results also suggest that both PVIs and sighted persons constructed accurate cognitive maps of the tactile environment on a behavioural level. However, early PVIs showed lower performance compared to late PVIs, suggesting an advantage of visual experience. Additionally, we reveal functional connectivity between areas that were involved in the navigation task during a subsequent resting-state block. This might suggest either visual imagination of stimuli during the preceding tasks, or cognitive processes related to our spatial navigation task, which possibly involve replay of stimulus-specific activity.

## Introduction

For successful navigation, we need a mental representation, or cognitive map, of our environment. Such a map contains spatial information, and spatial relationships such as distances between relevant locations. The human hippocampal formation stores such distances when the environment is explored using vision (Deuker, Bellmund, Navarro Schröder, & Doeller, 2016; Howard et al., 2014; Morgan, MacEvoy, Aguirre, & Epstein, 2011; Spiers & Barry, 2015). It is currently unclear whether this also happens when vision is reduced or not available, for instance in persons with a visual impairment (PVIs). In the present study, we investigated cognitive maps in the hippocampal formation based on non-visual information, whereby we focused on the haptic sensory modality. We aimed to assess whether the hippocampus and entorhinal cortex represent distances between locations on a small-scale tactile city-like map, in PVIs and sighted control participants.

Previous neuroimaging studies have provided evidence for the involvement of the hippocampus and entorhinal cortex in representing distances between relevant spatial locations (Howard et al., 2014; Spiers & Barry, 2015). During navigation, for instance, activity of the hippocampus correlates with path distance to the current goal (Howard et al., 2014; Spiers & Barry, 2015; Viard, Doeller, Hartley, Bird, & Burgess, 2011), and activity of the entorhinal cortex is related to Euclidean distance (Howard et al., 2014; Spiers & Barry, 2015). Furthermore, there is evidence that the hippocampus also stores distance information between relevant locations in a cognitive map (De Haas, Ottink, & Doeller, 2023; Deuker et al., 2016; Morgan et al., 2011). For instance, hippocampal pattern similarity between locations decreased with remembered distance between locations (Deuker et al., 2016). Moreover, Euclidean and path distances between relevant locations were represented and integrated by the hippocampus, as reflected in neural pattern similarity between locations (De Haas et al., 2023).

Involvement of the hippocampus and entorhinal cortex in representing spatial distances has been in navigation studies where participants rely on vision. In the current study, we aimed to investigate whether a cognitive map in the hippocampus-entorhinal cortex is also formed based on non-visual sensory information, in particular the haptic modality. Many studies have shown that persons with a visual impairment (PVIs) can form a cognitive map of a tactile environment, as assessed using behavioural tasks (e.g., Gagnon et al., 2012; Gaunet, Martinez, & Thinus-Blanc, 1997; Miao, Zeng, & Weber, 2017; Papadopoulos, Koustriava, & Kartasidou, 2012; Thinus-Blanc & Gaunet, 1997). Behavioural measures include for example estimating distances between relevant locations, and recalling specific locations. Both PVIs and sighted persons have been shown to form accurate mental representations of distances on a tactile map (Afonso et al., 2010; Blanco & Travieso, 2003; Ottink, Van Raalte, Doeller, Van der Geest, & Van Wezel, 2022a; ThinusBlanc & Gaunet, 1997) and of specific locations (Gaunet et al., 1997; Iachini, Ruggiero, & Ruotolo, 2014; Papadopoulos et al., 2012; ThinusBlanc & Gaunet, 1997). Some studies indicate similar distance representations in PVIs and sighted persons (Ottink et al., 2022a; Thinus-Blanc & Gaunet, 1997). There is also research, however, suggesting that especially PVIs who had their visual impairment since birth (early PVIs) have slightly worse mental representations of distances compared to PVIs who acquired their visual impairment at a later age (late PVIs) and sighted persons (Afonso et al., 2010; Blanco & Travieso, 2003). Considering mental representation of locations on a tactile map, some research indicates a lower accuracy in early PVIs (Gaunet et al., 1997; Iachini et al., 2014), however, with extensive training, mental representations of specific locations are similar in PVIs and sighted persons (Papadopoulos et al., 2012; ThinusBlanc & Gaunet, 1997).

In short, previous research suggests that PVIs as well as sighted persons can form a mental representation of a tactile map on a behavioural level (Ottink et al., 2022b). It is currently unclear whether learning an environment in a non-visual sensory modality also results in a neural map-like representation similarly as in the visual domain. In the present study, we aimed to address this issue by assessing the formation of a cognitive map in the hippocampus and entorhinal cortex based on haptic information. Only few studies have investigated neural activation in the context of exploration of a tactile map. For instance, PVIs showed increased activation of right hippocampus, parahippocampus, occipital cortex, fusiform gyrus, and precuneus during tactile maze solving, while sighted participants showed increased activation of the caudate nucleus, thalamus and precuneus (Gagnon et al., 2012). We also explored BOLD responses during tactile map navigation.

To investigate the formation of cognitive maps in the hippocampal formation, our participants performed a navigation task on a small-scale tactile city-like map, and learned five item locations in this environment. We used adaptation analysis to assess representations of distances between the five item locations in the hippocampus and entorhinal cortex. Adaptation analysis is based on the observation that when a stimulus is presented twice, the neural response to the second stimulus is reduced (adaptation effect; Barron, Garvert, & Behrens, 2016; Grill-Spector & Malach, 2004; Krekelberg, Boynton, & van Wezel, 2006). We predict that items close together in space would activate overlapping populations of spatially tuned cells in the hippocampal formation, such as place cells and grid cells, and therefore show an adaptation effect when subsequently presented (Barron et al., 2016; Garvert, Dolan, & Behrens, 2017). Therefore, when two items from the tactile map would be presented subsequently, we expected that the adaptation effect would scale with the distance between the two items on the learned map. Our task design did not allow to disentangle between Euclidean and path distance, therefore, we assessed representations of Euclidean distances as these are more indicative of a map-like representation than path distances (Eichenbaum, Dudchenko, Wood, Shapiro, & Tanila, 1999; Foo, Warren, Duchon, & Tarr, 2005; Schinazi, Thrash, & Chebat, 2016). To this end, our participants performed an item listening task before and after the navigation task, where only the five items from the tactile map were repeatedly presented in subsequent pairs. Additionally, they completed a distance estimation and an item location recall task to behaviourally assess cognitive map formation. We tested for differences between PVIs and sighted participants in behavioural performance as well as neural representations of distance.

Furthermore, we explored effects of navigational strategies on distance representations in the hippocampal formation. We assessed differences between allocentric and egocentric navigators (Astur, Purton, Zaniewski, Cimadevilla, & Markus, 2016; Burgess, Maguire, & O’Keefe, 2002; Iglói, Doeller, Berthoz, Rondi-Reig, & Burgess, 2010). These two strategies might lead to differential neural representations (Iglói et al. 2010; De Haas et al., 2023). Previous studies have demonstrated that egocentric navigators show representation and integration of distances in the right hippocampus, but allocentric navigators do not (De Haas et al., 2023). We distinguished between allocentric and egocentric navigators by categorising our participants into place learners and response learners, respectively, using an adapted T-maze task (Astur et al. 2016; De Haas et al., 2023). We created an auditory version of this task, to make it suitable for PVIs. We tested for differences between place and response learners in behavioural performance as well as neural representations of distance. To address general navigational abilities, and general navigational strategy use in daily life, participants furthermore filled out two questionnaires at the end of the experiment. The Santa Barbara Sense of Direction Scale (SBSOD; Hegarty, Richardson, Montello, Lovelace, & Subbiah, 2002) assessed navigational abilities and sense of direction, and the Wayfinding Strategy Scale (WSS; Lawton, 1994; Prestopnik & Roskos-Ewoldsen, 2000) indicates whether participants have a tendency to use a survey strategy (related to an allocentric perspective), or a route strategy (related to an egocentric perspective; Prestopnik & Roskos-Ewoldsen, 2000).

Additionally, we explored functional resting-state connectivity in PVIs and sighted participants, to assess possible cognitive processing related to haptic spatial navigation during a resting-state block after performing the navigation task in the MRI-scanner. In this analysis, correlations of time series of different brain regions are calculated, indicating the strength of functional connectivity between these regions. Early blind individuals have shown higher connectivity of memory regions to the visual cortex compared to sighted persons, suggesting stronger incorporation of the visual cortex in memory and attention to non-visual tasks (Burton, Snyder, & Raichle, 2014). The visual cortex may have furthermore deviated to processing of non-visual sensory information or even other cognitive functions in early PVIs (Yu et al., 2008). Moreover, early PVIs exhibited lower connectivity of visual cortex to other sensory areas compared to sighted persons (Burton et al., 2014; Wang et al., 2014). These alterations may reflect processes such as control of attention and suppression of distracting inter-sensory activity. This could possibly guide behaviour in early PVIs to adjust to their environment without the use of vision (Burton et al., 2014; Wang et al., 2014). These results have been found unrelated to a task. In the current experiment, we wanted to explore such connectivity in the context of spatial navigation, where both PVIs and sighted participants had to perform non-visual tasks. Analysing resting-state functional connectivity in this context may give more insight in for instance consolidation or post-encoding spatial processing of multimodal spatial information in PVIs and sighted participants. We speculate that this might also relate to whether and how they had constructed a cognitive map. Therefore, we analysed functional connectivity between the hippocampus, entorhinal cortex and visual cortex, to other sensory regions in PVIs and sighted participants during a subsequent resting-state block. We included the auditory cortex and sensorimotor cortex (hand region), as our navigation tasks involved auditory and tactile stimuli.

In short, our main aim was to investige representations of distances on a tactile city-like map in the hippocampus and entorhinal cortex, and whether these are different in PVIs and sighted participants. We furthermore addressed effects of navigational strategies on these neural representations. Lastly, we explored functional connectivity between areas involved in the navigation task and sensory areas in PVIs and sighted participants.

## Methods

### Tactile map

In our experiments, we used a small-scale tactile city-like map, of approximately 20 × 30 cm (Figure 1B), with five locations marked by half spheres on the middle of the streets. The streets had a width of 16 mm, and were 7 mm lower than the surroundings. We placed small bars like speed bumps around the intersections, such that participants would notice they had arrived on an intersection where they could change direction. Additionally, several tactile textures were detectable along the edges of the surroundings, to facilitate orientation.

**Figure 1.**
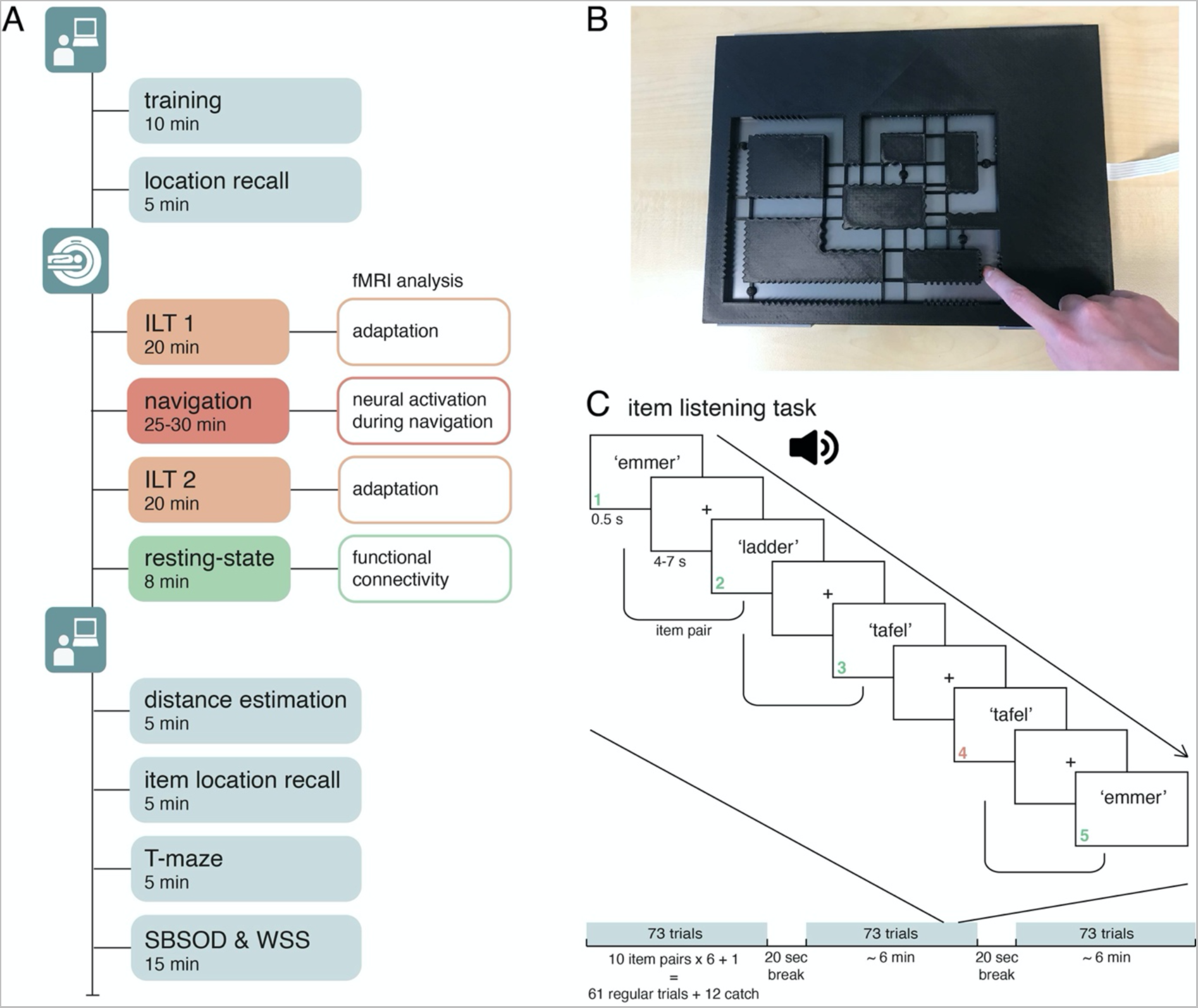
Overview of the experimental design, tactile map, and item listening tasks. **A**. Overview of the experimental design. The experiment starts with a behavioural session, which includes the training (10 min) and the location recall task (~5 min). The subsequent MRI-session includes ILT 1 (~20 min), the navigation task (~25 min), ILT 2 (~20 min), and a resting-state block (8 min). ILT 1 and 2 were analysed using adaptation analysis, to investigate neural representations of distances. MRI-data from the navigation task were analysed to assess neural activation during haptic navigation. Data from the resting-state block were analysed to investigate functional connectivity between memory and sensory regions. The second behavioural session included the distance estimation task (~5 min), item location recall task (~5 min), the T-maze task (~5 min), and the SBSOD and WSS questionnaires (~15 min). **B**. The tactile map on a touch panel. The streets were open, so the touch panel could track the position of the participants on the map. The five location markings are visible as circles on a bar on the middle of the streets. Small bars were placed around the intersections like speed bumps, such that participants would notice they had arrived on an intersection where they could change direction. Several tactile textures were detectable along the edges of the surroundings, to facilitate orientation. **C**. Overview of the item listening task. Examples of items that were repeatedly presented as spoken words are shown. Items were presented for 500 ms, and the ITI was either 4000, 5500, or 7000 ms. In this example, trial 1, 2, 3 and 5 are regular trials, and 4 is a catch trial, where the item is the same as in the preceding trial. Two items presented subsequently were considered an item pair, i.e., in this example, ‘emmer’ and ‘ladder’ (trial 1-2) are a pair, as well as ‘ladder’ and ‘tafel’ (trial 2-3), and ‘tafel’ and ‘emmer’ (trial 4-5). The order of the items was pseudorandomised, such that each item pair was presented an equal number of times. Each ILT was divided into 3 blocks of ~6 min. Each pair was presented 6 times per block (3 times in both directions). There were 10 item pairs, yielding 61 regular trials per block (60 items that formed the ‘second item of a pair’, plus 1 item at the start of each block to complete the 60 pairs). During each block, 12 randomly determined trials were followed by a catch trial, yielding 73 trials per block, and 219 in total per ILT. Between blocks, participants had a break of 20 seconds.

During the navigation task, the locations were associated with unique items, which were presented as a Dutch, spoken word. These items were chosen such that they appear approximately equally often in Dutch language, and were two-syllable words. This was determined using SUBTLEX-NL database (http://crr.ugent.be/isubtlex/).

Participants were instructed to navigate from item to item, using only their right index finger, without making any jumps. To ensure this, the experimenter monitored the participants during the tasks. Furthermore, the streets of the map were open, and the map was placed on a touchpanel. This allowed us to track when the participants arrived at a certain item location on the map.

### Experimental session overview

In our experiment, we wanted participants to form a cognitive map of a small-scale tactile city-like map. To this end, they performed a navigation task using the tactile map, and several additional spatial tasks. An overview is given in Figure 1A and below. All tasks are described in more detail later on in the Methods section.

At the start of the experiment, the researcher gave an overview of the session. After these general instructions, the participants performed a training session to familiarise themselves with the tactile map, and learn the five locations and the routes between them (Figure 1A). At this point, no items were associated with the locations yet. Following this training, they performed a location recall task where they had to indicate the five locations on a similar tactile map without location markers. During a subsequent MRI-session, participants performed the navigation task using the same map as the training (Figure 1A). At this point, unique items were associated with the five locations. By navigating from item to item, participants learned the five items locations and the routes between them, and built up a cognitive map of the tactile environment. Before and after the navigation task, we repeatedly presented the five items in subsequent pairs in item listening tasks (ILT 1 and ILT 2, respectively; Figure 1A), during an fMRI-session. This allowed us to analyse signatures of a cognitive map in the hippocampus and entorhinal cortex using adaptation analysis. By subtracting the adaptation effect across item pairs in ILT 1 from ILT 2, we captured the adaptation effect, thus the representation of distances between locations, caused by associating items to the particular locations on the tactile map during the navigation task. At the end of the MRI-session, if there was time, we scanned a resting-state block to analyse functional connectivity between regions that had been involved in the navigation task.

Following the fMRI session, we tested cognitive map formation behaviourally. Here, participants had to perform a distance estimation task where they had to estimate distances between all item pairs, and an item location recall task where they had to indicate the five item locations on a similar map without location markers (Figure 1A). Finally, they performed an auditory T-maze task to assess place and response learning, and filled out the Santa Barbara Sense of Direction Scale (SBSOD) and Wayfinding Strategy Scale (WSS) questionnaires to assess general navigation abilities and general use of navigation strategies (Figure 1A). At the end of the experimental session, the experimenter asked several questions about the visual impairment of PVIs, autonomous navigation experience, and experience in braille and in orientation and mobility (O&M) training.

During behavioural tasks, participants sat at a table, with the tactile map in front of them. Sighted participants and PVIs with some residual vision were blindfolded while performing these tasks. During the navigation task in the MRI-scanner, the tactile map was placed on their lower belly, in the same orientation as when they sat at a table. All participants reported that they experienced the map very similarly when sitting at a table and when lying in the MR-scanner. During tasks where participants received auditory instructions or stimuli, they wore headphones, or MR-compatible in-ear headphones during the MRI-session.

### Participants

In total, 47 people participated in our experiment, of which 22 were visually impaired (11 female; mean age = 45, SD = 14; see Table 1 for details), and 25 were sighted controls (12 female; mean age = 47, SD = 16) in our study. There was enough time in the MRI-session to finish the resting-state block for 37 of those participants (18 PVIs, 19 sighted controls). We included an additional group of 21 sighted participants for a short behavioural experiment, who only completed the T-maze task and the SBSOD and WSS questionnaires, to validate the auditory version of the T-maze task. Participants were recruited via advertisements distributed via client organisations and personal contacts of researchers and participants.

**Table 1.** Details of participants with a visual impairment: age, gender, cause of visual impairment, age of onset of visual impairment, autonomous navigation, O&M training experience, and braille experience.

| Age | Gender | Cause of visual impairment | Age of onset | Autonomous navigation | O&M training experience | Braille experience |
| --- | --- | --- | --- | --- | --- | --- |
| 35 | Male | No eyes | 0 | Often | Yes | Yes |
| 29 | Female | Leber Congenital Amaurosis | 0 | Often | Yes | Yes |
| 18 | Male | Leber Congenital Amaurosis | 0 | Never | Yes | Yes |
| 26 | Female | Leber Congenital Amaurosis | 0 | Often | Yes | Yes |
| 45 | Female | Retinitis pigmentosa | 0 | Often | Yes | Yes |
| 55 | Male | Retinal damage at birth | 0 | Often | Yes | Yes |
| 63 | Male | Retrolental fibroplasia | 0 | Regularly | Yes | Yes |
| 57 | Female | Congenital cataract | 0 | Often | Yes | Yes |
| 32 | Male | Glaucoma, aniridia | 0 | Often | Yes | Yes |
| 56 | Male | Retinitis pigmentosa | 0 | Regularly | Yes | Yes |
| 50 | Female | Premature birth | 0 | Often | Yes | Yes |
| 26 | Female | Retinitis pigmentosa | 13 | Often | Yes | Yes |
| 30 | Female | Aniridia | 18 | Often | Yes | Yes |
| 59 | Male | Optical nerve damage | 21 | Never | Yes | Yes |
| 47 | Female | Stargardt disease | 23 | Often | Yes | Yes |
| 42 | Male | Proliferative retinopathy | 26 | Often | Yes | Yes |
| 31 | Male | Usher syndrome | 28 | Often | Yes | No |
| 57 | Female | Dominant cystoid macular dystrophy | 42 | Often | No | No |
| 56 | Female | Familial exudative vitreoretinopathy | 45 | Often | Yes | Yes |
| 64 | Female | Cone-rod dystrophy | 50 | Often | No | Yes |
| 62 | Male | Retinitis pigmentosa, Choroideremia | 55 | Often | No | Hardly |
| 61 | Male | Eye infection | 57 | Often | Yes | No |

An inclusion criterium for participants with a visual impairment was that they are not able to use or read visual street maps at all without support. Additionally, we only included PVIs whose cause of visual impairment is not located in the brain (Table 1). None of the PVIs and sighted participants had a sensory or cognitive impairment other than visual for the PVIs. A visual impairment corrected to normal by glasses or contacts was allowed for the sighted participants. All participants were right-handed and Dutch-speaking.

The study was ethically approved by the local ethical committee via an amendment (CMO Regio Arnhem-Nijmegen, The Netherlands, nb. 2014/288). All participants gave informed consent before the start of the experiment (written by sighted participants, and recorded on video by PVIs which are stored separately from the rest of the data).

Sighted participants and PVIs with any residual vision were blindfolded during all spatial tasks when sitting at a table. They did not wear a blindfold in the MRI-scanner, as they could not see the tactile map or anything else related to the spatial tasks while lying down.

We excluded one PVI from analyses, based on movement during the second item listening task. One sighted participant was excluded from analyses because that participant could not finish the navigation task because of time issues. Both these participants were only included in the resting-state analyses.

### Spatial tasks

#### Training

The aim of the training (Figure 1A) was for the participants to learn the layout of the map, including the five locations and the routes between them. At this time, no items were associated with the locations. Participants could familiarise themselves with the map and the various components such as the location markers, speed bumps around intersections, and tactile textures in the surrounding walls (Figure 1B).

Participants started at one of the locations, where they were directed by the experimenter. They could then freely navigate the map using their right index finger for 10 minutes. Participants were instructed to only follow the streets and not make any shortcuts. Each time they reached a location, this was registered through the touch panel, by a script programmed in MATLAB (MATLAB and Statistics Toolbox Release 2017b, The MathWorks, Inc., Natick, Massachusetts, United States). Participants were monitored by the experimenter. If there was a location that they could not find, they were directed there by the experimenter, to make sure all participants had experienced all five locations at least several times before moving on to the next task.

#### Navigation task

During the navigation task (Figure 1A), unique items were associated with the five locations on the map. The goal of this task was for the participants to learn these item location by navigating from item to item, and further learn the map layout and routes between the item locations. They started at one of the locations, as indicated by the experimenter. At the start of a trial, participants received instruction about which item they had to find. When they reached this target item location, they received feedback about the item that was located there. They were then instructed about the next item they had to find, and so on. The order of the target item locations was pseudorandomised such that the route between each item pair was navigated 6 times (3 times in both directions). There were 5 items, therefore 10 pairs, leading to a total of 60 trials. The item pairs (routes) were equally distributed across the task, to ensure that a particular route is not only experienced early or late in the task.

Similar to the training, participants were instructed to only navigate using their right index finger, and only follow the streets. When they arrived at the target item location, this was registered through the touch panel. The task was programmed in MATLAB (MATLAB and Statistics Toolbox Release 2017b, The MathWorks, Inc., Natick, Massachusetts, United States). The average completion time of this task was 26.19 minutes for PVIs (SD = 4.25), and 28.99 minutes for sighted participants (SD = 4.98 min).

#### Distance estimation task

After the MRI-session, participants had to estimate relative Euclidean and path distances of the shortest route between each item pair. Their estimations had to lie on a scale from 0 to 100, where 0 would mean two items are placed on the same location, and 100 would be the distance between the items that are furthest apart. Distances between the other item pairs had to be linearly scaled to this. Participants first had to estimate all Euclidean distances, followed by all path distances. The item pairs were randomised for each participant. The estimations were given verbally by the participants, and recorded by the experimenter in a data file.

#### Location and item location recall tasks

Participants had to recall the five locations on the tactile map in two similar tasks. They performed a location recall task after the training, when no items were associated with the locations yet. After the navigation task, where items had been associated with the locations, participants had to recall these item locations.

##### Location recall task

After the training, participants had to recall the five locations on the map (Figure 1A). They had to point out the remembered locations on the tactile map without location markers. Participants could freely choose the order of locations. They navigated on the map, and notified the experimenter when they arrived at a remembered location. The coordinates where then registered by a script programmed in MATLAB (MATLAB and Statistics Toolbox Release 2017b, The MathWorks, Inc., Natick, Massachusetts, United States). After this task, if the experimenter noticed that a participant did not remember one of the locations or deviated a lot from the correct locations, they guided them past all locations once more before proceeding to the next task.

##### Item location recall task

Similarly to the location recall task, participants had to point out the item locations on the map without location markers during the item location recall task (Figure 1). They received instructions about which item they had to indicate, and notified the experimenter when they arrived at its remembered location on the tactile map. These coordinates were registered by a script programmed in MATLAB (MATLAB and Statistics Toolbox Release 2017b, The MathWorks, Inc., Natick, Massachusetts, United States). The order of the items was randomised for each participant.

### Navigational ability tasks

#### T-maze

To categorise participants into place learners (allocentric navigators) and response learners (egocentric navigators), we built an auditory version of a T-maze task, adapted from a visual version (Astur et al., 2016; De Haas et al., 2023). In this task, participants were placed in a virtual auditory environment, and stood at the bottom of a T-shaped intersection. Several auditory landmarks were present in the environment (such as chirping birds, people talking in the distance, and a fountain), distinguishable at the left or right side through headphones. Participants were instructed to find a reward in the left or right arm of the T, which they could choose pressing the left or right button on a keyboard. They had to do this for 10 trials, and the reward was always in the same location in the environment. For 2 of the 10 trials, the T and thus the participant was rotated 180°, but the landmarks remained at their location (probe trials). If people were place learners, they would notice that the auditory landmarks had rotated, and follow this (i.e., turn left during probe trials when they had to turn right to find the reward in regular trials, or vice versa; they learn the place of the reward). If people were response learners, they would not notice the rotation, and they would keep taking the same turn as they had to take to find the reward in regular trials (they learn their response). During probe trials, both responses (i.e., turning left and right) were rewarded. The 10^th^ trial always was a probe trial, and the first probe trial was random the 6^th^ or 8^th^ trial. The task was programmed in MATLAB (MATLAB and Statistics Toolbox Release 2017b, The MathWorks, Inc., Natick, Massachusetts, United States).

When a participant chose the correct arm during both probe trials, we categorised them as place learner. When a participant chose the incorrect arm during both probe trials, we categorised them as response learner. If a participants once chose the correct and once the incorrect arm, they were categorised as mixed. In addition to the sighted control participants who completed the whole experiment, we conducted the T-maze task with an additional group of 21 sighted participants. They also filled out the SBSOD and WSS questionnaires, to validate the auditory version of the T-maze task.

Across the PVIs, 12 participants were categorised as place learners, 7 as response learners, and 2 as mixed. Across sighted who completed the whole experiment, 11 were categorised as place learners, 11 as response learners, and 2 as mixed. Of additional sighted participants, 6 were categorised as place learner, 13 as response learner, and 2 as mixed. Therefore, across all 45 sighted participants who performed the T-maze task, 17 were place learners, 24 response learners, and 4 were mixed.

#### Questionnaires

At the end of the experiment, participants filled out two questionnaires, the Santa Barbara Sense of Direction (SBSOD; Hegarty et al. 2002) to assess general navigational abilities and sense of direction, and the Wayfinding Strategy Scale (WSS; Lawton 1994; Prestopnik and Roskos-Ewoldsen 2000) to assess general navigation strategy use. The questionnaires had been translated to Dutch. Both questionnaires yielded a score between 0 and 7 for each participant. A higher score indicated better general navigational abilities for the SBSOD, and tendency to use a survey strategy for the WSS. A lower score indicated worse general navigation abilities for the SBSOD, and a tendency to use a route strategy for the WSS.

The mean SBSOD-score of PVIs was 4.581 (SD = 0.968), and the mean SBSOD-score of sighted participants (who finished the whole experiment) was 4.847 (SD = 1.135). This suggest a fairly good self-report of general navigation abilities in both groups. The mean WSS-score of PVIs was 3.191 (SD = 0.636), and 3.313 for sighted participants (SD = 0.674). This indicates no clear self-reported tendency to use a route or survey strategy in both groups.

### Behavioural analyses

We behaviourally assessed cognitive map formation using distance estimation and item location recall tasks. To analyse performance on the distance estimation task, we correlated estimated Euclidean and path distances with the correct distances, using Pearson correlation. For further analyses, we Fisher transformed these correlation coefficients. We tested whether correlation coefficients were significantly different from zero using 1-sample t-tests. Additionally, we tested for differences between PVIs and sighted participants, between place and response learners, and between Euclidean and path distance, using 2-sample t-tests.

To analyse the location recall task, we first determined the order in which the locations most likely were pointed out, because participants freely chose the order. Then, for both the location recall as well as the item location recall task, we calculated the Euclidean distance from the placed location to the correct location. We normalised this distance error by dividing it by the maximum distance error possible from that location. We then calculated the mean across locations, yielding one error score for each participant for both tasks. We tested whether participants significantly improved from the location recall to the item location recall task, using a nonparametric Wilcoxon signed rank test for paired samples (because the error scores are restricted between 0 and 1). Additionally, we tested for differences between PVIs and sighted participants and between place and response learners using nonparametric Wilcoxon rank sum tests for independent samples.

Furthermore, we correlated self-reported navigational abilities (SBSOD) and general wayfinding strategy use (WSS) with the scores on the distance estimation (using Pearson correlation) and item location recall task (using Spearman correlation). In addition, we tested for differences between place and response learners on the distance estimation task using a 2-sample t-test, and on the item location recall task using a Wilcoxon rank sum test. We also tested whether place and response learners and PVIs and sighted participants report different navigational abilities (SBSOD) or general wayfinding strategy use (WSS) using 2-sample t-tests.

### Item listening tasks

Before and after the navigation task, participants performed an item listening task (ILT 1 and ILT 2 respectively; Figure 1A), which allowed us to analyse signatures of a cognitive map in the hippocampus and entorhinal cortex using adaptation analysis. During these ILTs, only the items that were associated with the locations on the tactile map, were repeatedly presented as spoken words during an fMRI session (Figure 1C). Participants were instructed to actively listen to these items, and to perform an orthogonal 1-back task in order to keep their attention. Each time they heard an item, they had to press one of two buttons. When they heard the same item twice in a row (catch trials; Figure 1C) they had to press one button, and the other button when an item was different from the previous trial (regular trial; Figure 1C). Buttons were always pressed using the right index (left) and middle finger (right). Left or right buttons for regular or catch trials were counterbalanced across participants. Performance on the 1-back task showed a ceiling effect in both ILTs (ILT 1: mean = 91.5%, SD = 11.7; ILT 2: mean = 97.7%, SD = 3.7).

The items were presented in pseudorandomised order, but constant across the two ILTs of the same participant. Each pair of items (one item presented following another item) was presented an equal number of times. For example, when item A is followed by B, then followed by C, then A-B is a pair, and B-C is a pair, and so on (Figure 1C). Presentation of the items in pairs allowed us to later perform adaptation analysis. Each ILT was divided into 3 blocks, and each pair was presented 6 times per block (3 times in both directions, e.g., A-B and B-A). There were 10 item pairs, yielding 61 regular trials per block (60 items that formed the ‘second item of a pair’, plus 1 item at the start of each block to complete the 60 pairs, see Figure 1C), therefore 183 regular trials per ILT. During each block, 12 randomly determined trials were followed by a catch trial, where the same item was presented as during the regular trial. Therefore, the total number of trials was 219 for each ILT. Between blocks, participants had a break of 20 seconds, where they received feedback about their performance and a reminder of when to press which button. The item presentation was approximately 500 ms. The intertrial interval (ITI) was either 4000, 5500, or 7000 ms. The order of ITIs was pseudorandomised such that they were distributed equally across item pairs and blocks.

### fMRI analysis

#### fMRI acquisition

During ILT 1, the navigation task and ILT 2, functional T2*-weighted and anatomical images were acquired on a Magnetom Skyra 3 Tesla MRI-scanner (Siemens, Erlangen, Germany). Functional images were acquired using a multiband sequence (acceleration factor 4, 68 slices, multi-slice mode interleaved, TR = 1500 ms, TE = 33.4 ms, flip angle = 75 deg, FOV = 213 × 213 × 136 mm, isotropic voxel size 2 mm). Anatomical images of the brain were obtained using a T1 sequence (MPRAGE; TR = 2300 ms, TE = 3.03 ms, flip angle = 8 deg, FOV = 256 × 256 × 192 mm, isotropic voxel size 1 mm). After the navigation task, we furthermore acquired a gradient fieldmap scan to be able to correct for distortions, using a multiband sequence (TR = 510 ms, TE 1 = 2.80 ms, TE 2 = 5.26 ms, flip angle = 60 deg, isotropic voxel size 2 mm).

#### fMRI preprocessing

We preprocessed images of the three functional sessions (ILT 1, the navigation task and ILT 2) using the FSL toolbox (version 6.0.3, http://fsl.fmrib.ox.ac.uk/fsl/fslwiki/). We applied motion correction (three translation and three rotation estimations), a high-pass filter (cut-off 100s), and B0 unwarping using the acquired fieldmap scan. We brain-extracted the anatomical scans using BET, and downsampled them to the voxel size of the functional images (isotropic 2 mm). Subsequently, we linearly registered the functional images to the downsampled anatomical images, so we would have the same reference space in the functional sessions. Before moving on to the adaptation analysis, we excluded participants who gave no appropriate responses during the ILTs (the minimum criterium was that they had pressed both buttons), and based on motion (if more than 10% of volumes exceeded 4 mm movement). We excluded one PVI from analysis, based on movement during ILT 2.

#### First-level adaptation analysis

To measure representations of distances in the hippocampus and entorhinal cortex, we conducted adaptation analyses on the fMRI data recorded during the ILTs. This type of analysis is based on observations that when a stimulus is presented twice, neurons that respond to the first stimulus, show a reduced response during the second stimulus (Krekelberg et al., 2006). When stimuli (items) are associated to particular locations in space, we assumed that if they are close together in that space, their presentation activates an overlapping population of neurons (Barron et al., 2016). Therefore, when these two items are presented subsequently, the fMRI BOLD response to the second item would be reduced compared to the first. If two items are further apart in space, their presentation would activate a less overlapping population in neurons, leading to a lower reduction in BOLD response to the second item compared to two items close together in space (Barron et al., 2016; Garvert et al., 2017). We therefore predicted that across the whole ILT, the strength of the BOLD response to the second item of a pair would related to the distance between the associated locations on the tactile map.

We analysed the adaptation effect before (ILT 1) and after (ILT 2) the navigation task (Figure 1A) using general linear models. To obtain the effect caused by associating items to locations on the map, we subtracted the effect in ILT 1 from the effect in ILT 2. Because our task design did not allow to disentangle between Euclidean and path distance, we assessed representations of Euclidean distances only, as these are more indicative of a map-like representation than path distances (Eichenbaum et al., 1999; Foo et al., 2005). We used the Euclidean distance as predictions for the adaptation effect. We applied ROI-analyses to assess adaptation effects in the hippocampus and entorhinal cortex.

##### General linear model

We modeled the adaptation effect in a general linear model (GLM). Here, we modeled the onset and duration of each item presentation during the ILT in one regressor. In this ‘adaptation regressor’, the value of the parametric modulator for each item was set to the distance of that item to the preceding item, because we predicted that across the whole ILT, the response to an item would relate to the distance to the preceding item. We set up a GLM with the Euclidean distance prediction, for ILT 1 and ILT 2. In both GLMs, we also modeled the presentation of the five items in itself (onset and duration) in five regressors. Furthermore, we set up additional regressors for catch trials, button presses (by the index and middle finger in two regressors, using a stick function), the start of the task (from start time of scanning until the first item presentation), the end of the task (from end of the last item presentation until end of scanning), breaks between blocks (in one regressor, from the end of the last item of a block until the first item onset after the break), and six additional movement parameters (estimated during preprocessing). To extract the adaptation effect, we set a contrast that only included the adaptation regressor. This resulted in one contrast image for each participant, for each ILT.

##### ROI analysis

To analyse representations of distances in the left and right hippocampus and left and right entorhinal cortex, we performed ROI analyses using masks of these regions. Hippocampal masks were obtained from the Harvard-Oxford subcortical structural atlas (https://fsl.fmrib.ox.ac.uk/fsl/fslwiki/Atlases), and entorhinal cortex masks from a previous study (Navarro Schröder, Haak, Jimenez, Beckmann, & Doeller, 2015). We created shared ROI masks, which only included voxels that were grey matter voxels in all participants. These shared ROI masks were in MNI space, and we also registered the contrast images of each participant to MNI space. Subsequently, we calculated the mean parameter estimate of these contrast images within each shared ROI mask for each participant, for both ILT 1 and ILT 2. To get the effect caused by associating items to the locations on the tactile map, we subtracted the mean parameter estimate of each participant of ILT 1 (before the navigation task) from ILT 2 (after the navigation task). This yielded one value per ROI per participant.

#### Group-level adaptation analysis

We performed group level analysis on the ROI analysis in each participant. Furthermore, we also looked for effect outside the hippocampus and entorhinal cortex in a whole-brain analysis.

##### ROI analysis

For each ROI, we tested whether the adaptation effect (the mean parameter estimate across the ROI) was significantly different from zero across all participants using a 1-sample t-test. We furthermore tested for differences between PVIs and sighted participants, and between place and response learners, using 2-sample t-tests.

##### Whole-brain analysis

After registering the contrast images from participant to MNI space and spatial smoothing (6 mm full width at half maximum), we subtracted those of ILT 1 from ILT 2 for each participant. Subsequently, we performed a permutation test on the resulting contrast images using FSL randomise (10000 permutations). We used 1-sample permutation tests to analyse the adaptation effect across all participants, and 2-sample permutation tests for effects between PVIs and sighted participants. We applied threshold-free cluster enhancement, and family-wise error correction for multiple comparisons.

#### First-level neural activation during navigation analysis

To measure brain activity while exploring the tactile map, we analysed the fMRI data recorded during the navigation task (Figure 1A). We modeled this activation in each participant in a GLM. We set up a regressor that modeled all navigation periods, so when the participants navigated on the tactile map using their right index finger. This period was determined for each trial, as from the end of the instructions at the start of the trial, until the target location of that trial was reached. We also included six movement parameters (estimated during preprocessing). We set up a contrast that only included the navigation regressor. We registered the resulting contrast image of each participant to MNI space.

#### Group-level navigation analysis

We analysed whole-brain group-level activation during navigation periods of the navigation task using permutation tests. First, we created a shared whole-brain mask that only included voxels in all participants. We then analysed activation within the PVI and sighted groups, using permutation tests with small volume correction within the shared whole-brain mask (with 5000 permutations, threshold-free cluster enhancement, and family-wise error correction). To this end, we created a design matrix including all participants, and set contrasts to investigate group means for the PVI and for the sighted group. We furthermore performed 2-sample permutation tests to assess differences between the PVIs and sighted participants.

### Resting-state session

A subset of participants, when there was enough time, additionally completed a resting-state block of 8 minutes at the end of the MRI-session (Figure 1A). In total, 37 participants completed this session, of which 18 PVIs and 19 sighted participants. During this block, the MRI-room was darkened, to create similar conditions for PVIs and sighted participants. They were furthermore instructed to close their eyes, to not think of anything in particular, and not fall asleep.

#### Resting-state fMRI acquisition

Functional T2*-images were acquired on a Magnetom Skyra 3 Tesla MRI-scanner (Siemens, Erlangen, Germany). Functional images were acquired using a multiband sequence (acceleration factor 6, 66 slices, multi-slice mode interleaved, TR = 1000 ms, TE = 35.2 ms, flip angle = 60 deg, FOV = 213 × 213 × 132 mm, isotropic voxel size 2 mm). After the resting state block, a gradient fieldmap scan was acquired, using a multiband sequence (TR = 500 ms, TE 1 = 2.80 ms, TE 2 = 5.26 ms, flip angle = 60 deg, isotropic voxel size 2 mm). For resting state connectivity analyses, we used the same anatomical T1 scan as for the adaptation analysis.

#### Resting-state fMRI preprocessing

We brain-extracted the anatomical scans using the FSL Anatomical Processing Script (version 6.0.3, http://fsl.fmrib.ox.ac.uk/fsl/fslwiki/fsl_anat). We used this as reference image for preprocessing of the functional resting-state run, using the FSL toolbox (version 6.0.3, http://fsl.fmrib.ox.ac.uk/fsl/fslwiki/). We applied motion correction, no temporal filtering, 5 mm full width at half maximum (FWHM) spatial smoothing, B0 unwarping using the acquired fieldmap scan, and we deleted the first 5 volumes to allow for the signals reaching equilibrium. Additionally, we performed non-aggressive denoising, removing only the variance uniquely associated with the noise components, using the FSL-based toolbox ICA-AROMA (ICA-based automatic removal of motion artifacts; Pruim, Mennes, van Rooij, et al. 2015; Pruim, Mennes, Buitelaar, et al. 2015). In addition to removal of motion components, we also regressed out effects of white matter and CSF, by extracting their mean timeseries.

#### Seed-based connectivity analysis

We performed seed-based connectivity analysis (SBCA) on the resting state functional run. Here, we analysed functional connectivity between three structurally defined seed ROIs (hippocampus, entorhinal cortex and V1), and several target regions that are involved in the navigation task that participants performed earlier during the fMRI session (auditory cortex, sensorimotor cortex of the hand, occipital cortex, and hippocampus-entorhinal cortex). The same hippocampal and entorhinal cortex masks were used as for the adaptation ROI analyses. These masks were registered to functional space for each participant. Furthermore, we determined a V1-mask for each participant using the FreeSurfer software (version 7.2.0, https://surfer.nmr.mgh.harvard.edu/fswiki). An auditory cortex mask was determined using the HCP-MMP1 Cortex Labels atlas (including early and association auditory cortex; https://fsl.fmrib.ox.ac.uk/fsl/fslwiki/Atlases). A mask for the sensorimotor cortex of the hand was determined from a cortical parcellation atlas (Cortical Area Parcellation from Resting-State Correlations; Gordon et al., 2016). Furthermore, a occipital cortex mask was created using the Harvard-Oxford cortical structural atlas (including occipital pole, lateral occipital cortex inferior and superior, and occipital fusiform gyrus; https://fsl.fmrib.ox.ac.uk/fsl/fslwiki/Atlases). The target ROI masks were in MNI space.

##### First-level SBCA

For each participant, we determined the functional connectivity from the seed ROIs to the rest of the brain. We analysed the signal correlation in each voxel to the mean time courses in the seed ROI masks using the FSL SBCA function (https://fsl.fmrib.ox.ac.uk/fsl/fslwiki; fsl_sbca), and a mask of the seed ROI and of whole-brain in functional space. We registered the resulting correlation image of each participant to MNI space.

##### Group-level SBCA

We analysed the group-level connectivity from the seed ROIs to whole-brain and the target regions. First, we analysed connectivity from the seed ROIs to whole-brain across all participants. Therefore, we created a shared whole-brain mask that only contained voxels in all participants. We then performed a 1-sample nonparametric permutation test on the participant correlation images in the shared whole-brain mask, with 5000 permutations, threshold-free cluster enhancement, and family-wise error correction for multiple comparisons. For descriptive purposes, we also explored connectivity in the hypothesised target ROIs (auditory, sensorimotor, occipital and hippocampus-entorhinal cortices; small-volume corrected, and using a p-threshold of 0.017 for cluster detection to correct for multiple ROIs), within the PVI and sighted group using permutation tests.

To then explore differences between PVIs and sighted participants in the target ROIs, we determined the target ROIs, for each seed ROI, using the auditory, sensorimotor, occipital and hippocampus-entorhinal cortex masks, in which we only included voxels that showed significant whole-brain connectivity to the seed ROI across all participants. Subsequently, we analysed differences in connectivity between the seed ROIs to the target ROIs between the PVI and sighted groups, using 2-sample permutation tests with small volume correction within the target ROIs (with 5000 permutations, threshold-free cluster enhancement, and family-wise error correction). Here, we created a design matrix including all participants, and set contrasts to investigate differences between the PVIs and sighted participants, as well as differences between early and late PVIs.

## Results

### Behavioural signatures of a cognitive map

We measured cognitive map formation behaviourally using a distance estimation, and an item location recall task. Both the PVIs and sighted participants performed well on the distance estimation task, as indicated by high correlation of their estimated distances with the actual distances (Figure 2A). The median correlation of estimated and actual Euclidean distance was r = 0.834 (p < 0.001) for PVIs, and r = 0.869 (p < 0.001) for sighted participants. The median correlation of estimated path distance was r = 0.827 (p < 0.001) for PVIs, and r = 0.792 (p < 0.001) for sighted participants (Figure 2A). After Fisher transformation of the correlation coefficients, we found no differences between PVIs and sighted participants for Euclidean (p = 0.617, T_43_ = −0.504) or path distance (p = 0.364, T_43_ = −0.918). We did, however, find differences between PVIs who had their visual impairment since birth (early PVIs) and who acquired it at a later age (late PVIs; Figure 2C). On Euclidean distance estimation, early PVIs perform significantly worse compared to late PVIs (p < 0.001, T_19_ = −4.179) and sighted participants (p < 0.05, T_33_ = −2.637). Early PVIs also perform worse than late PVIs (p < 0.05, T_19_ = −2.810) and sighted (p < 0.05, T_33_ = −2.399) participants on path distance estimation. We found no differences between late PVIs and sighted participants on Euclidean (p = 0.087, T_32_ = 1.764) or path distance estimation (p = 0.266, T_32_ = 1.132).

**Figure 2.**
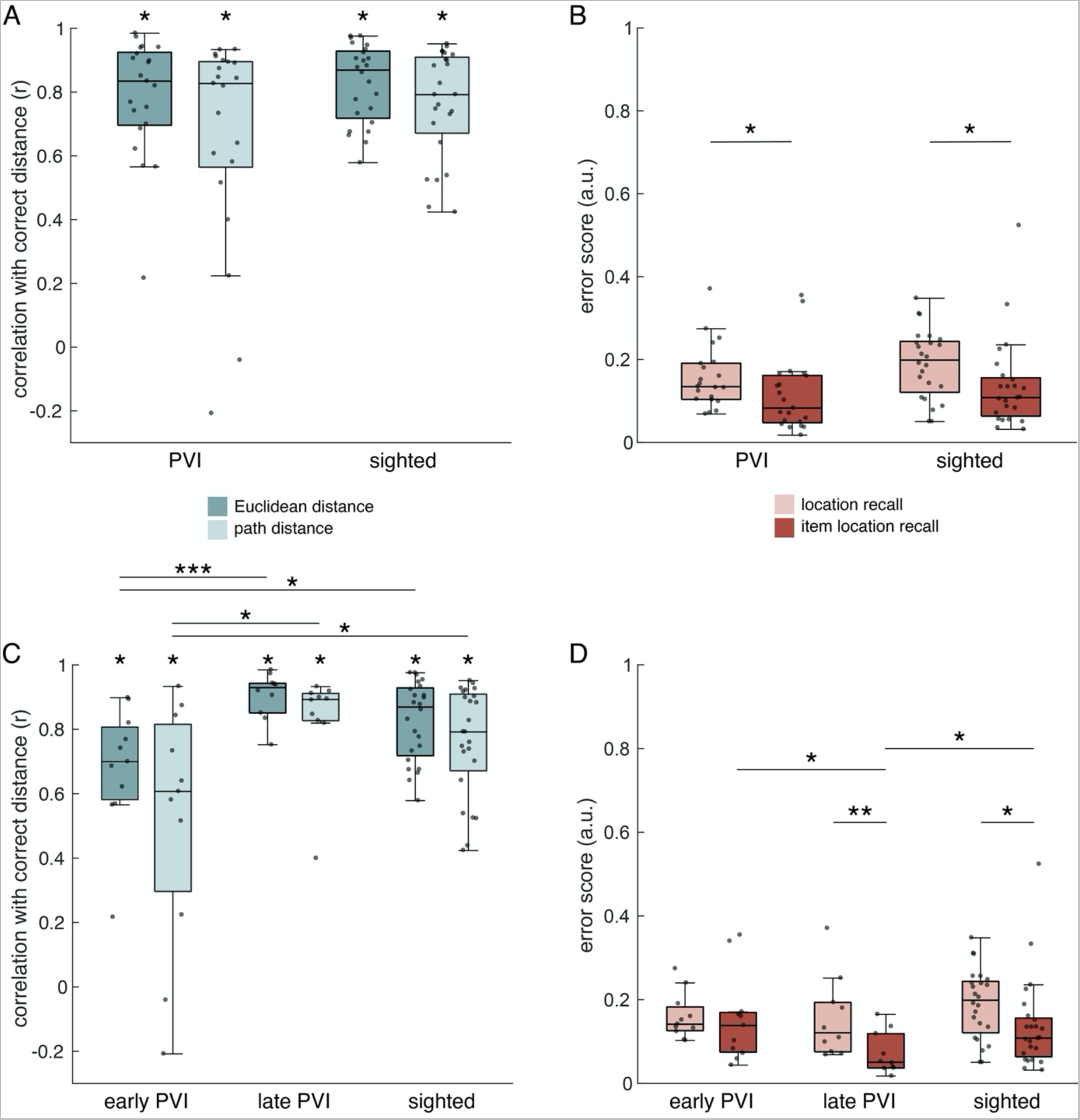
Behavioural performance on the distance estimation, location recall, and item location recall tasks. **A**. Correlation of estimated distance with actual distance of the distance estimation task, showing results of Euclidean and path distance estimations, for PVIs and sighted participants. **B**. Error scores on the location recall and item location recall tasks, showing results for PVIs and sighted participants. **C**. Correlation of estimated distance with actual distance of the distance estimation task, showing results of Euclidean and path distance estimations, for early PVIs, late PVIs, and sighted participants. **D**. Error scores on the location recall and item location recall tasks, showing results for early PVIs, late PVIs, and sighted participants. Boxplots indicate median performance, and 25^th^ and 75^th^ percentiles. Grey dots indicate individual data points. *p < 0.05, **p < 0.01, ***p < 0.001

On the item location recall task, both PVIs and sighted participants obtain fairly low error scores (Figure 2B, dark boxes), indicating good memory of item locations on the tactile map. Both groups show significant improvement from the location to the item location recall task (PVIs: p = 0.042, W = 165; Sighted: p = 0.037, W = 168; Figure 2B). This may suggest that associating items to the locations, as well as simply encountering them more often in the environment, significantly improved knowledge about the five locations. There was no difference between the PVIs and sighted participants on the location recall (p = 0.155, W = 420) as well as the item location recall task (p = 0.433, W = 448). On the item location recall task, however, we found that early PVIs had a significantly higher error score than late PVIs (p = 0.018, W = 76), as well as sighted compared to late PVIs (p = 0.043, W = 121), however, early PVIs and sighted participants still scored fairly well. There was no difference between early PVIs and sighted participants (p = 0.511, W = 217). Furthermore, early PVIs showed no significant improvement from location recall to item location recall task (p = 0.831, W = 10), however, late PVIs did (p = 0.002, W = 55; Figure 2D). Taken together, performance on the distance estimation as well as the item location recall task indicate accurate cognitive map formation on a behavioural level by PVIs and sighted participants.

We furthermore correlated the performance on the distance estimation and item location recall tasks with scores on the SBSOD and WSS questionnaires. We found a significant relationship between self-reported navigational abilities (as measured with the SBSOD) and path distance estimation in PVIs (r = 0.513, p = 0.017), but not sighted participants (r = 0.047, p = 0.828). Furthermore, we discovered a negative correlation between SBSOD scores and error scores on the item location recall task in PVIs (r = −0.441, p = 0.045), but not in sighted participants (r = 0.172, p = 0.422). No significant relationships of behavioural performance with self-reported navigational strategy use (as measured with the WSS) were revealed.

We also tested for differential behavioural performance by place and response learners. We found a marginally significant difference on the item location recall task in sighted participants where place learners had a higher error score than response learners (p = 0.049, W = 96), but not in PVIs (p = 0.432, W = 80). In addition, we analysed place and response learning in an supplementary sample of sighted participants, to validate the auditory version of the T-maze task. Across all sighted participants, including the supplementary sample (n = 46), we found that place learners score higher on the SBSOD, indicating higher self-reported general navigation abilities compared to response learners (place learners: mean = 5.086; response learners: mean = 4.440; p = 0.028, T_40_ = 2.273). We do not find this effect in PVIs, nor in the smaller sample of sighted participants who completed the whole experiment. We might therefore need a large sample size to find such an effect. However, the relationship between place and response learning and SBSOD-scores suggests validity of our own developed auditory version of the T-maze task.

We could not test for effects of experience in autonomous navigation, O&M training or braille reading, as almost all participants (19 out of 22) had high competence in these areas. We did test for effects of age, however, we did not discover a significant correlation of age with performance on either the distance estimation or item location recall task.

### Neural signatures of distance representations

To analyse cognitive map formation in the hippocampal formation, we performed adaptation analysis on the fMRI data recorded during the ILTs (Figure 1A). We used Euclidean distance as predictions for the adaptation effect, and conducted ROI analyses on the left and right hippocampus, and left and right entorhinal cortex. Across all participants, we found no significant effect in the left (p = 0.426, T_44_ = −0.804) or right (p = 0.372, T_44_ = −0.903) hippocampus (Figure 3A). However, in the left entorhinal cortex, we observed a significant effect (p = 0.034, T_44_ =−2.194), but not in the right entorhinal cortex (p = 0.749, T_44_ = −0.322; Figure 3B). We found a negative effect in the left entorhinal cortex (Figure 3B), which means a higher adaptation effect for item locations that are further apart in space, and a lower adaptation effect for item locations closer together in space.

**Figure 3.**
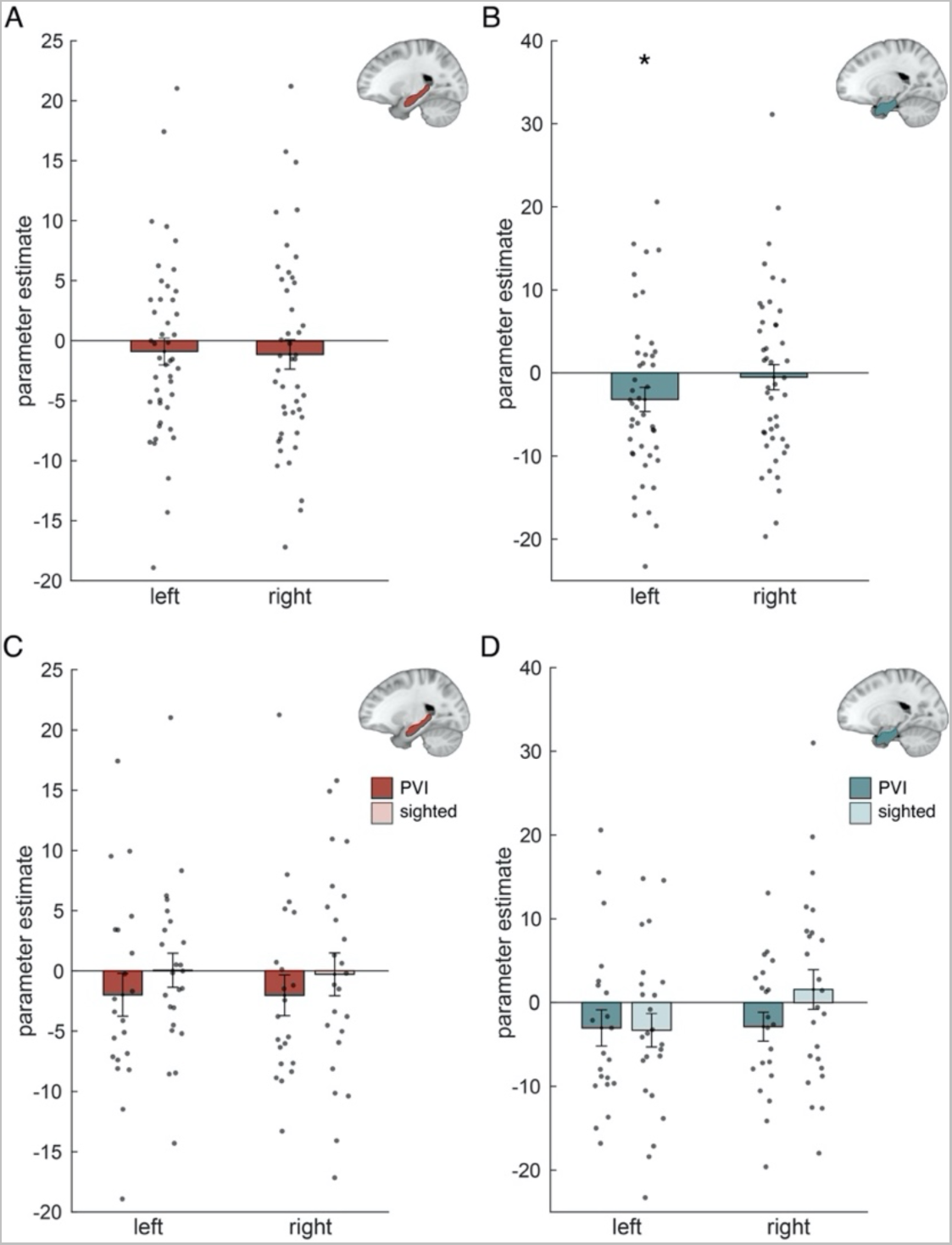
Adaptation effect of Euclidean distance **A**. Adaptation results in the left and right hippocampus across all participants. **B**. Adaptation results in the left and right entorhinal cortex across all participants. **C**. Adaptation results in the left and right hippocampus in PVIs and sighted participants. **D**. Adaptation results in the left and right entorhinal cortex in PVIs and sighted participants. Bars indicate mean ± SEM. Grey dots are individual data points. *p < 0.05

We did not find significant effects when testing for PVIs and sighted participants separately (Figure 3C and D), possibly due to the low power in the smaller group sizes. Likewise, we found no differences between PVIs and sighted participants, nor between early and late PVIs, or between place and response learners across all participants. Furthermore, we correlated the effect in the left entorhinal cortex with performance on the distance estimation and item location recall tasks, and with self-reported general navigation abilities (SBSOD questionnaire) and navigation strategy use (WSS questionnaire). We found one marginally significant positive correlation of error scores on the item location recall task with Euclidean distance representation in sighted (r = 0.413, p = 0.046), but not PVIs. No other relations were discovered.

To explore representations of distance outside the hippocampal formation, we additionally performed a whole-brain analysis. A one-sample permutation test on a whole-brain grey matter mask revealed no significant effect, corrected for multiple comparisons. A two-sample permutation test also revealed no differences between PVIs and sighted participants. We show the uncorrected effects in Figure 4. As a posthoc analysis, we checked the location of the peak in the left entorhinal cortex using the whole-brain t-stats image across all participants. The peak voxel was located in the medial part of the left entorhinal cortex (T = 3.176, MNI-coordinates X −17, Y −4, Z −27; entorhinal cortex division based on Schröder et al., 2015).

**Figure 4.**
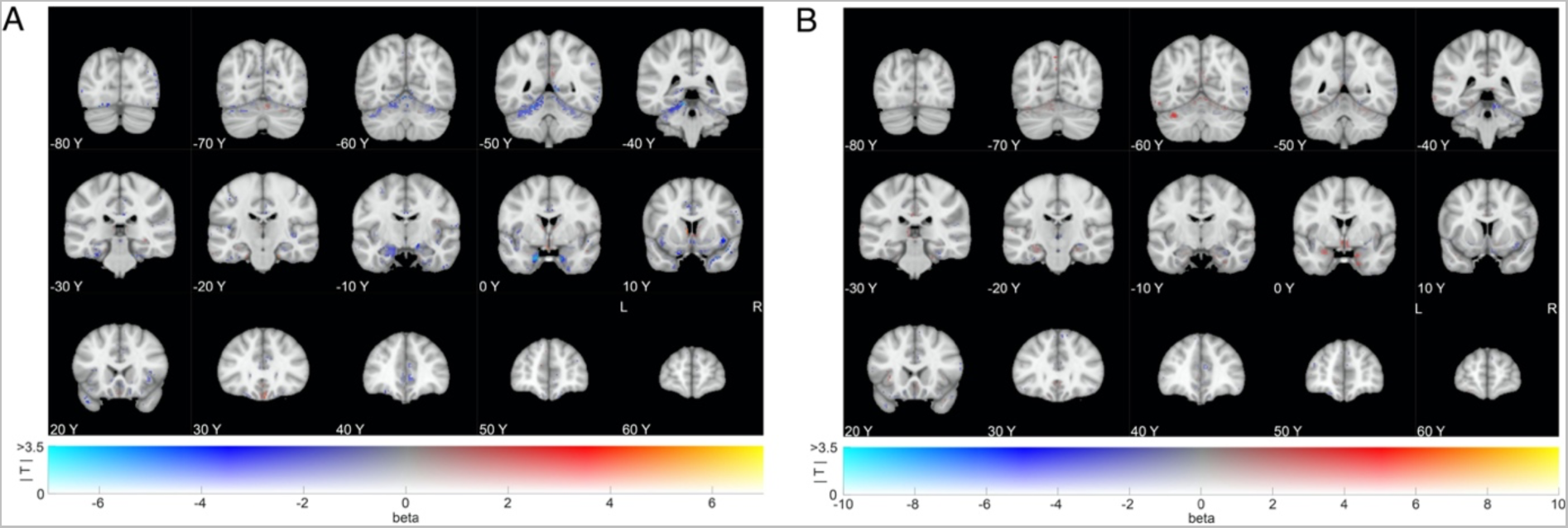
Whole-brain adaptation effects. **A**. Whole-brain adaptation results across all participants. No effects survived whole-brain correction. **B**. Whole-brain adaptation results for differences between PVIs and sighted participants. No effects survived whole-brain correction. Images were created using dual-coding (Allen, Erhardt, & Calhoun, 2012; Zandbelt, 2017). Mean beta coefficients are indicated by color, and t-stats by opacity. Coordinates are in MNI-space.

### Neural activation during the navigation task

We analysed neural activation during the navigation periods of the navigation task, in PVIs and sighted participants. Across all PVIs, we found one significant cluster in the precuneus. In sighted participants, we found significant activation of the precuneus, posterior cingulate cortex, left somatosensory cortex, left primary motor cortex, supplementary motor area, and right occipital cortex. Activation of the left somatosensory and motor cortices corresponds to the right hand, which participants used to navigate on the tactile map. Activation of the occipital cortex might relate to visual imagery of the tactile map. Furthermore, the precuneus is generally involved in motor imagery, and directing attention for movements in space (Cavanna & Trimble, 2006). Activation of the precuneus and occipital cortex during tactile maze solving has been found in earlier research (Gagnon et al., 2012). The cingulate cortex is generally involved in learning and memory (Stanislav, Alexander, Maria, Evgenia, & Boris, 2013), and the supplementary motor area in planning movements and coordinating sequences of movements (Lee & Quessy, 2003). Significant clusters are reported in Table 2. We found no differences between PVIs and sighted participants that survived whole-brain correction when testing across all PVIs.

**Table 2.** Significant clusters of activation during navigation periods of the navigation task within the early and late PVI and sighted group, using a p-threshold of 0.05. Cluster size, maximum t-value, and MNI-coordinates of peak voxels are reported.

| Region | Early PVI |  |  |  |  | Late PVI |  |  |  |  | Sighted |  |  |  |  |
| --- | --- | --- | --- | --- | --- | --- | --- | --- | --- | --- | --- | --- | --- | --- | --- |
|  | Cluster size | Max t-value | MNI |  |  | Cluster size | Max t-value | MNI |  |  | Cluster size | Max t-value | MNI |  |  |
|  |  |  | X | Y | Z |  |  | X | Y | Z |  |  | X | Y | Z |
| Precuneus, posterior cingulate, left somatosensory, left primary motor, supplementary motor and right occipital cortex | - | - | - | - | - | 2682<br>43 | 6.39<br>4.68 | -12<br>-26 | -6<br>14 | 44<br>54 | 2392 | 5.94 | -14 | -30 | 56 |

Furthermore, we checked for differences between early and late PVIs. We found that early PVIs did not significantly activate regions during navigation periods that survived whole-brain correction. Late PVIs, however, activated similar regions as sighted participants did: precuneus, posterior cingulate cortex, left somatosensory cortex, left primary motor cortex, and right occipital cortex (Table 2). In these regions, we also found significant stronger activation by late PVIs and sighted participants compared to early PVIs. We did not find differential activation between late PVIs and sighted participants.

### Functional connectivity between navigation-related areas

We analysed functional connectivity during a resting-state block, between three structurally defined seed ROIs (hippocampus, entorhinal cortex and V1), and several target regions that were involved during the navigation task (auditory cortex, sensorimotor cortex of the hand, occipital cortex, and hippocampus-entorhinal cortex). We investigated functional connectivity across all participants using seed-based connectivity analysis, and tested for differences between PVIs and sighted participants. For descriptive purposes, we also explored functional connectivity within the PVI and sighted groups. Significant clusters from this exploratory analysis and their peak voxels are reported in Table 3.

**Table 3.**
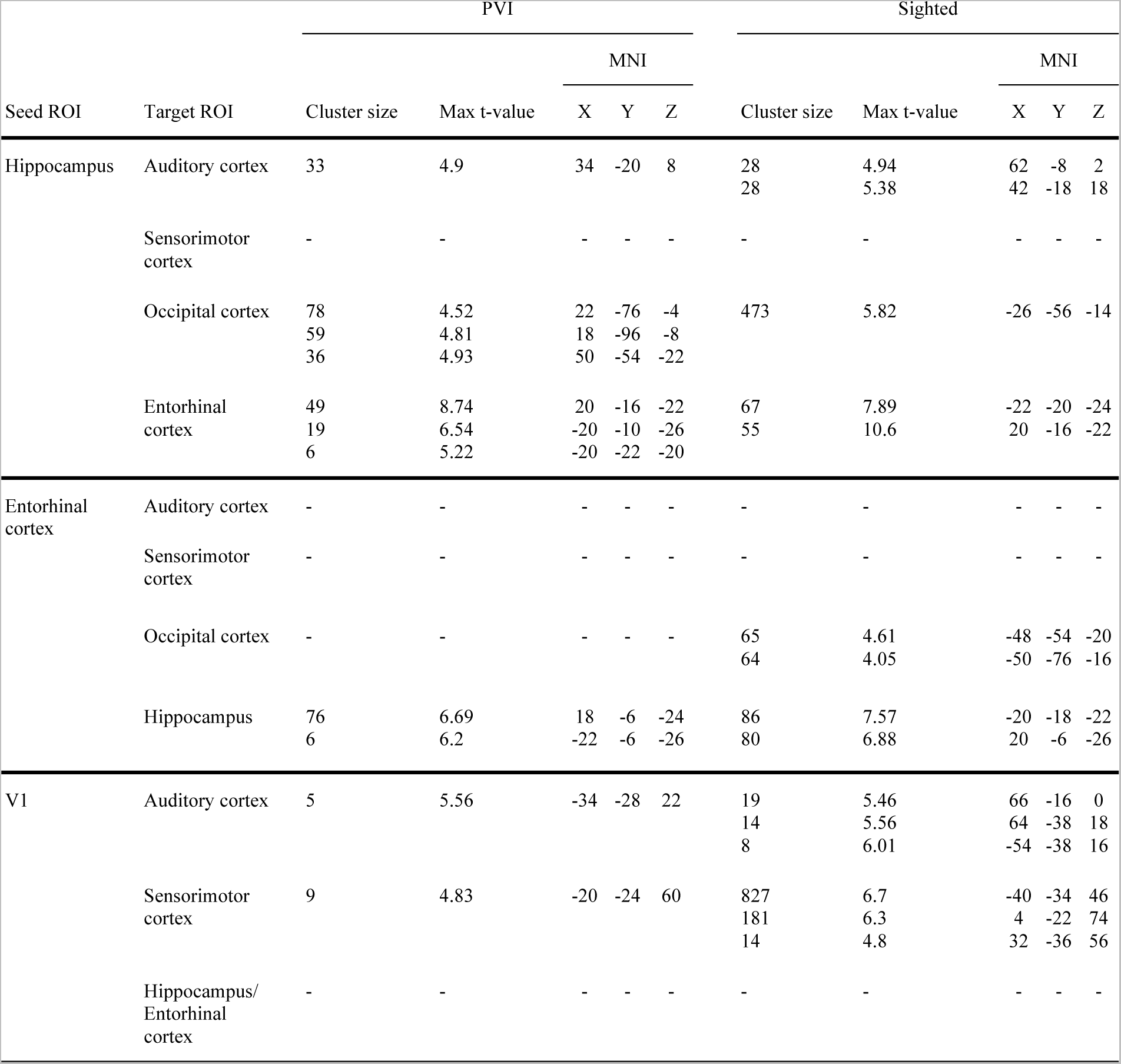
Significant clusters of functional connectivity between seed ROIs and hypothesised target ROIs, within the PVI and sighted group, using a p-threshold of 0.017 to correct for multiple ROIs. Cluster size for clusters larger than 5 voxels, maximum t-value, and MNI-coordinates of peak voxels are reported.

#### Connectivity from the hippocampus

Across all participants, we observed significant connectivity from the hippocampus across a large portion of the brain, including to all hypothesised target ROIs, after whole-brain correction. We did not reveal significant differences between PVIs and sighted participants, nor when separating PVIs into early and late PVIs.

For descriptive purposes, we also explored functional connectivity to the target ROIs (small-volume corrected) within the PVI and sighted groups. There is significant connectivity in PVIs and sighted participants from the hippocampus to the right auditory cortex. Furthermore, we found significant clusters in the occipital cortex for both sighted participants and PVIs (in the lateral occipital and fusiform gyrus). We found no connectivity to the sensorimotor cortex of the hand that survived multiple comparisons correction for either PVIs or sighted participants. We did find significant clusters in the entorhinal cortex for both groups.

#### Connectivity from the entorhinal cortex

Across all participants, we found significant connectivity from the entorhinal cortex to the hippocampus, left lateral occipital cortex, and right auditory cortex after whole-brain correction. We found no significant differences between PVIs and sighted participants, or significant distinctions when separating PVIs into early and late PVIs.

For descriptive purposes, we also explored functional connectivity to the target ROIs (small-volume corrected) within the PVI and sighted groups. The entorhinal cortex does not exhibit functional connectivity to the auditory or sensorimotor cortex (hand region) in both groups. We did find a significant cluster in the occipital cortex in sighted participants (left lateral inferior occipital cortex), but not PVIs. Furthermore, the entorhinal cortex shows connectivity to the left and right hippocampus in both PVIs and sighted participants.

#### Connectivity from the primary visual cortex

Across all participants, we found significant connectivity from the primary visual cortex (V1) across a large portion of the brain, including to all hypothesised target ROIs, after whole-brain correction. We did not reveal significant differences between PVIs and sighted participants, or significant distinctions when separating PVIs into early and late PVIs.

For descriptive purposes, we also explored functional connectivity to the target ROIs (small-volume corrected) within the PVI and sighted groups. We found significant connectivity from V1 to the left auditory cortex in PVIs, and to the left and right auditory cortex in sighted participants. Furthermore, we found significant clusters in the left sensorimotor cortex of the hand in PVIs, and in the left and right sensorimotor cortex in sighted participants. Here, sighted participants show much larger clusters compared to PVIs. V1, however, does not show significant connectivity to the hippocampus-entorhinal cortex that survived multiple comparisons correction in PVIs or sighted participants.

## Discussion

The aim of the current study was to investigate behavioural and neural signatures of cognitive map formation based on haptic information, in visually impaired and sighted participants. Our data indicates that both groups can construct accurate mental representations of item locations on a tactile map and distances between them. These behaviourally measured cognitive maps were somewhat more accurate in late PVIs compared to early PVIs. Furthermore, we found evidence of distance representations in the left entorhinal cortex. Here, we did not discover differences between PVIs and sighted participants. Importantly, our data indicates preserved construction of cognitive maps in the hippocampal formation when an environment is presented in a non-visual sensory modality, pointing towards the formation of modality-independent maps in the brain.

### Behavioural signatures of a cognitive map

We measured cognitive map formation on a behavioural level using distance estimation and item location recall tasks. Our results indicate accurate formation of mental representations in both PVIs and sighted participants. Across all PVIs, we found no differences in distance estimation or item location recall performance between PVIs and sighted participants. This is similar to our previous research (Ottink et al., 2022a). When testing for differences between early and late PVIs, a comparison that we could not make in the previous study, however, we show that early PVIs experienced somewhat more difficulties estimating distances and recalling item locations compared to late PVIs and sighted, although their performance is still fairly good. We speculate that the lower performance might relate to the significantly lower activation of spatial cognition-related areas during the navigation task in early PVIs compared to late PVIs and sighted participants. Additionally, sighted participants formed slightly less accurate mental representations of item locations compared to late PVIs, but not of distances. An advantage of visual experience in incorporating spatial information into a mental representation has been suggested by earlier research, for both distances (Afonso et al., 2010; Blanco & Travieso, 2003; Cattaneo et al., 2008) and specific locations (Gaunet et al., 1997; Iachini et al., 2014). Visual experience may promote visual imagery, which might in turn support retrieving information from cognitive maps (Gaunet et al., 1997). In addition, in our tasks, late

PVIs might have been able to use their visual experience in combination with experience in using non-visual sensory modalities for spatial tasks, to perform superior to both early PVIs and sighter participants. Nevertheless, differences between early and late PVIs may diminish with extensive training on the tactile map, as has been previously suggested in multiple studies (Ottink et al., 2022b; Papadopoulos et al., 2012; ThinusBlanc & Gaunet, 1997; Ungar, 2000). Taken together, our data shows that both PVIs and sighted participants formed accurate cognitive maps of the tactile map on a behavioural level. The precision, however, is slightly lower in early PVIs compared to late PVIs.

### Neural signatures of a cognitive map

We investigated neural signatures of cognitive maps by analysing representations of distances between relevant locations on the tactile map in the hippocampus and entorhinal cortex. To this end, we performed adaptation analysis on the fMRI data recorded during ILT 1 and ILT 2. We expected that the adaptation effect across a whole ILT would relate to the distance between the presented items. We subtracted the adaptation effect in ILT 1 from ILT 2, to obtain the effect caused by associating items to the specific locations on the tactile map.

We found a significant effect in the left entorhinal cortex when testing across all participants. Our analysis revealed no difference between PVIs and sighted participants. This suggests that the ability to map space in the hippocampal formation may be preserved in persons to whom visual information is reduced or not available. This is in line with many behavioural findings showing that PVIs can construct cognitive maps (e.g., Gagnon et al., 2012; Gaunet et al., 1997; Miao et al., 2017; Papadopoulos et al., 2012; Thinus-Blanc & Gaunet, 1997). Our behavioural data, however, shows lower performance by early compared to late PVIs. This difference is not reflected in the adaptation effect, possibly because of low power when comparing neural data within the PVI group. We did, however, find significant lower activation of areas related to spatial cognition during the navigation task in early PVIs compared to late PVIs and to sighted participants. Importantly, our data suggests that a space learned through a non-visual sensory modality leads to a neural representation similar to a map based on visual information. This contributes to ideas of modality independent coding of space. This concept proposes that information from different sensory modalities are combined into one multimodal representation rather than a distinct representation from each modality (Loomis, Klatzky, & Giudice, 2013; Tcheang, Bulthoff, & Burgess, 2011; Wolbers, Klatzky, Loomis, Wutte, & Giudice, 2011).

The involvement of the entorhinal cortex in spatial distance coding has been suggested in multiple lines of research. For instance, place cells have not only been discovered in the hippocampus, but also upstream, in the entorhinal cortex (Fyhn, Molden, Witter, Moser, & Moser, 2004), and the entorhinal cortex provides information about multisensory environmental cues to the hippocampus (Witter, Doan, Jacobsen, Nilssen, & Ohara, 2017). Furthermore, entorhinal cortex activity has been shown to be related to Euclidean distances (Howard et al., 2014; Spiers & Barry, 2015; Spiers & Maguire, 2007). Our data supports this evidence. Specifically, entorhinal cortex activity has been shown to be related to the Euclidean distance to the current goal (Howard et al., 2014; Spiers & Maguire, 2007). During the ILTs in our experiment, the items that were related to specific locations on the tactile map, were subsequently presented. When for instance item B is presented following item A, the entorhinal cortex may process item B as the current goal from A, resulting in the observed effect (Howard et al., 2014; Spiers & Maguire, 2007). Furthermore, posthoc analysis shows that the peak adaptation effect in the left entorhinal cortex is located in the medial part of the EC, which is more associated with spatial information, while the lateral part of the EC is more affiliated with multisensory and item-related information (Navarro Schröder et al., 2015). This further supports the involvement of the entorhinal cortex in spatial distance coding.

Nevertheless, we expected an adaptation effect, therefore a positive relationship between neural activity and distance between item locations. Our data, however, shows a negative relationship. Such an association has been shown in earlier research, where activity in the entorhinal cortex correlated negatively with distance to the current goal (Spiers & Maguire, 2007). It might therefore be that our observed effect is not necessarily an adaptation effect (based on the assumption that locations close in space activate an overlapping population of neurons), but rather activity of the entorhinal cortex in relation to the distance to a new goal location, made visible because of the design of our ILTs. This still points towards the formation of a cognitive map, because during the ILTs, the items were presented unrelated to their location. Therefore, their location information has to had been stored in the brain. This storage, however, might have not been in the entorhinal cortex itself, but perhaps in the hippocampus, which communicated this information to the entorhinal cortex. The effect in the entorhinal cortex might have emerged over the hippocampus because of the multisensory nature of our navigation task. Another important thing to note here, is that distance-related activity has been previously observed in the right entorhinal cortex (Howard et al., 2014; Spiers & Maguire, 2007), while we show an effect in the left entorhinal cortex. However, we have shown distance coding in the left instead of right hippocampus in previous work (De Haas et al., 2023). We do not find evidence for such storage of distances in the hippocampus in our study. Because we do find distance-related coding in the entorhinal cortex, we speculate that our study design or our analysis methods might not be sufficient to detect representations of distances in the hippocampus. We speculate that it could be the case, for instance, that locations on the small-scale tactile map are not distinct enough. They might be all classified as close together by the hippocampus, in comparison to locations in a large-scale real-world environment, which is the scale our brain’s navigation system is calibrated to.

Taken together, we found neural signatures of cognitive map formation based on haptic information, in the left entorhinal cortex. Distance coding in the hippocampus based on haptic information, and potential neural differences between early and late PVIs have to be examined further with higher power. Nevertheless, importantly, our data shows the first neural evidence that the ability to map space in the hippocampal formation is preserved when an environment is presented in a non-visual sensory modality. This points towards the formation of modality-independent maps in the brain.

### Navigational abilities and strategies

We assessed the role of navigational abilities and strategies using an auditory version of a T-maze task and the Santa Barbara Sense of Direction Scale (SBSOD) and Wayfinding Strategy Scale (WSS) questionnaires. To test the validity of the auditory version of the T-maze, we performed this task with an additional sample of 21 sighted participants. When testing across all sighted participants (n = 46), we found that place learners have higher self-reported general navigation abilities compared to response learners. This indicates that also the auditory version of this simple behavioural task (Astur et al., 2016; De Haas et al., 2023) can divide participants into good and worse navigators. We only find this relationship between the T-maze strategy and SBSOD-scores in the large sample of sighted participants, not in the group of sighted participants who finished the whole experiment, or in the PVI group. We therefore might need a large sample size to detect this effect.

Nevertheless, when testing across all participants who finished the whole experiment (including PVIs and sighted, n = 45), we do not reveal a difference between place and response learners in SBSOD-score. There might be differences between PVIs and sighted persons that diminish the effect when taken together. For instance, we found that place learners perform somewhat worse on the item location recall task compared to response learners, in sighted but not in PVIs. This difference between place and response learners might arise because of the design of the item location recall task. To perform well on this task, it might be sufficient to associate a landmark such as the tactile textures or a specific turn on the map, to each location, and recall this during the item location recall task. Here, the fact that the tactile map is always perceived from the same direction may also play a role. People who navigate mostly based on such associations could be response learners, and might benefit on this particular task compared to place learners. However, this has to be particularly investigated. Moreover, we do not find differences between place and response learners on other behavioural tasks or in the fMRI adaptation results. Since we needed a larger sample size to show the relationship with the SBSOD-scores, this could also be the case for the other measures. Furthermore, from the questionnaires it was revealed that our participants are generally good navigators. The difference between our categorised ‘good’ and ‘worse’ navigators may not be large enough to distinguish between them on a behavioural or neural level.

We furthermore assessed whether there are relationships between scores on the SBSOD and WSS and other behavioural tasks. We show a positive relationship between self-reported navigational abilities (as measured using the SBSOD), and accuracy on path distance estimation and item location recall in PVIs, but not in sighted participants. Such a relationship has been found in sighted participants based on visual information in our previous work (De Haas et al., 2023). Given that the PVIs in the current study showed a similar correlation between task performance and self-reported abilities, this may be related to the different sensory modalities. PVIs may be more accustomed to using haptic information for spatial tasks compared to sighted, resulting in the demonstrated correlations. Sighted participants are less familiar with haptic spatial processing, but do exhibit similar correlations when the spatial tasks are based on visual information (De Haas et al., 2023). They do, however, perform similarly well on the spatial tasks based on haptic information as PVIs in the current study. Therefore, it could also be an issue of power, as the sighted sample size is higher in the previous study (De Haas et al., 2023), and the proposed relationships are not shown in PVIs or sighted in a study with smaller sample (Ottink et al., 2022a).

Moreover, we found no differences between place and response learners in distance representations in the hippocampus or entorhinal cortex across all participants. We speculate that this might for instance be related to the way of navigation during our experiment. While exploring a small-scale tactile map, people might not engage their usual navigation strategies as they would in a real-world environment, because the method of navigation is incredibly dissimilar. On a tactile map, people encounter all spatial information in close proximity to their own body. One could argue that it is therefore processed egocentrically, however, because the tactile map easily gives an overview of the environment, one could suggests it is processed allocentrically. This discrepancy could diminish distinctions in processing of spatial information between place and response learners. Another point to consider, is that there could be differences in place and response learners between PVIs and sighted participants, that reduce the effect when taken together. Our sample size is not sufficient to test for neural differences between place and response learners within the PVI and sighted groups. Besides, differences between place and response learners can establish in neural signatures in the hippocampus (De Haas et al., 2023; Iglói et al., 2010, 2015), however, we do not find distances representations in that region in the first place. It might additionally be the case that our participants were not distinct enough in their strategy use to give rise to a neural effect. This is also reflected in the similar scores on the Wayfinding Strategy Scale across participants. Besides, successful navigation may be related to effective switching between the two strategies rather than always employing one over the other (Gagnon, Kupers, Schneider, & Ptito, 2010; Kupers, Chebat, Madsen, Paulson, & Ptito, 2010), and egocentric and allocentric representations exist in parallel in the brain (Burgess, 2006). Even when participants employ only one strategy during the T-maze task, they might swith during an actual navigation task, especially because our T-maze and navigation task are presented in a different modality, diminishing the distinction between place and response learning.

### Functional connectivity between memory and sensory areas

In addition to behavioural and neural signatures of cognitive maps, we have explored functional connectivity in the context of a spatial navigation task, an approach not implemented with PVIs before. We analysed functional connectivity between regions implicated in this navigation task across all participants during a subsequent resting-state block. We did not discover any significant differences in connectivity between the groups.

First, we analysed connectivity from the hippocampus to sensory cortices and the entorhinal cortex. We found significant connectivity between the hippocampus and the auditory cortex across all participants. This could be related to items that were auditorily presented during the navigation task and the ILTs. We furthermore found connectivity that could be related to tactile stimulation by the map, or to replay of the tactile stimulation during navigation (Deuker et al., 2017), as significant connectivity between hippocampus and sensorimotor cortex of the hand was revealed. Important to note, however, is that the connectivity from the hippocampus is widespread across the brain when testing across all participants. Regarding specific activity related to tactile map stimulation, we would expect more specific connectivity to the left sensorimotor cortex (corresponding to the right hand), however, we found connectivity to the left and right cortex. Moreover, when exploring connectivity in PVIs and sighted participants separately, the connectivity to the sensorimotor cortex diminishes. We did discover significant connectivity to the occipital cortex across all participants. From data recorded during the navigation task, we also found significant activation of the occipital cortex while navigating on the tactile map. We speculate that this might be associated with visual imagination of the spatial task or as replay of processes that happened during the spatial task (Deuker et al., 2017; Zhang, Deuker, & Axmacher, 2017). We furthermore revealed significant connectivity between the hippocampus and entorhinal cortex, which is expected as the entorhinal cortex is the main input and output structure of the hippocampus (Witter et al., 2017).

As expected, we also found a significant cluster in the hippocampus from the entorhinal cortex seed across all participants. This cluster was located in the anterior portion of the hippocampus. Therefore, this connectivity might not be necessarily related to spatial, but rather to the processing or reactivation of recent memories (Dandolo & Schwabe, 2018), which might relate to our tasks. Furthermore, we also found connectivity to the auditory and occipital cortices across all participants. Similar to the hippocampus, this connectivity might be associated with the items that were auditorily presented during the ILT and navigation task, and visual imagination or replay. Nevertheless, this effect diminishes when exploring connectivity in PVI and sighted groups separately.

Finally, we investigated connectivity from the primary visual cortex (V1) to the auditory and sensorimotor cortices and to the hippocampal-entorhinal region. Our data revealed significant functional connectivity to the auditory and sensorimotor cortex, across all participants. Previous literature has suggested lower resting-state connectivity between V1 and other sensory cortices in early PVIs compared to sighted participants (Burton et al., 2014). The ‘cross-modal’ hypothesis, however, predicts stronger functional connectivity between the visual and other sensory cortices in PVIs compared to sighted persons (Burton et al., 2014; Wittenberg, Werhahn, Wassermann, Herscovitch, & Cohen, 2004). Nevertheless, both these findings are not supported by our data, as we do not reveal differences between PVIs and sighted participants, also not when splitting the PVIs into early and late PVIs. This discrepancy might arise because of the specific involvement of the auditory and sensorimotor cortices in our spatial navigation task, and possibly because of visual imagination of the auditory items and tactile map. In addition, we found significant resting-state connectivity between V1 and the hippocampal-entorhinal region across all participants. Important to note is, however, is that the connectivity from V1 is widespread across the brain when testing across all participants. When exploring connectivity in PVIs and sighted participants separately, the connectivity to the hippocampal-entorhinal region diminishes. Previous research indicated increased connectivity between V1 and the hippocampus in late PVIs compared to sighted participants (Wen et al., 2018). However, this is not replicated in our study, as we do not find any differences between early or late PVIs and sighted participants.

In short, our data shows functional connectivity between memory and sensory areas that have been involved in the spatial navigation task during a subsequent resting-state block. We found no differences between PVIs and sighted participants. This lack of difference is also reflected in the neural representations of distances in the hippocampal formation in the two groups, as measured earlier in the experiment. The resting-state connectivity might suggest visual imagination of stimuli during the preceding tasks, or post-encoding cognitive processes related to our spatial navigation task, which possibly involve replay of stimulus-specific activity (Deuker et al., 2017; Zhang et al., 2017).

## Conclusions

Importantly, our data shows the first neural evidence that the ability to map space in the hippocampal formation is preserved when an environment is presented in a non-visual sensory modality. This points towards the formation of modality-independent maps in the brain. We revealed neural representations of distances on a tactile map in the left entorhinal cortex across all participants, and did not find differences between PVIs and sighted participants. On a behavioural level, early PVIs perform slightly worse compared to late PVIs, however, the results still indicate accurate cognitive map formation. We speculate that this might relate to the significantly lower activation of spatial cognition-related areas during the navigation task in early PVIs compared to late PVIs and sighted participants. Finally, our results show functional resting-state connectivity between memory and sensory areas that have been involved in the spatial navigation task in PVIs and sighted participants. This might suggest visual imagination of stimuli during the preceding tasks, or post-encoding processing, which possibly involve replay of stimulus-specific activity.

## References

Afonso, A., Blum, A., Katz, B. F. G., Tarroux, P., Borst, G., & Denis, M. (2010). Structural properties of spatial representations in blind people: Scanning images constructed from haptic exploration or from locomotion in a 3-D audio virtual environment. Memory & Cognition, 38(5), 591–604.

Allen, E. A., Erhardt, E. B., & Calhoun, V. D. (2012). Data visualization in the neurosciences: overcoming the curse of dimensionality. Neuron, 74(4), 603–608.

Astur, R. S., Purton, A. J., Zaniewski, M. J., Cimadevilla, J., & Markus, E. J. (2016). Human sex differences in solving a virtual navigation problem. Behavioural Brain Research, 308, 236–243.

Barron, H. C., Garvert, M. M., & Behrens, T. E. J. (2016). Repetition suppression: A means to index neural representations using BOLD? Philosophical Transactions of the Royal Society B: Biological Sciences, 371(1705).

Blanco, F., & Travieso, D. (2003). Haptic Exploration and Mental Estimation of Distances on a Fictitious Island: From Mind’s Eye to Mind’s Hand. Journal OfVisual Impairment & Blindness, (May 2003), 298–300.

Burgess, N. (2006). Spatial memory: how egocentric and allocentric combine. Trends in Cognitive Sciences, 10(12), 551–557.

Burgess, N., Maguire, E. A., & O’Keefe, J. (2002). The Human Hippocampus and Spatial and Episodic Memory. Neuron, 35(4), 625–641.

Burton, H., Snyder, A. Z., & Raichle, M. E. (2014). Resting state functional connectivity in early blind humans. Frontiers in Systems Neuroscience, 8(1 APR), 1–13.

Cattaneo, Z., Vecchi, T., Cornoldi, C., Mammarella, I., Bonino, D., Ricciardi, E., & Pietrini, P. (2008). Imagery and spatial processes in blindness and visual impairment. Neuroscience & Biobehavioral Reviews, 32(8), 1346–1360.

Cavanna, A. E., & Trimble, M. R. (2006). The precuneus: A review of its functional anatomy and behavioural correlates. Brain, 129(3), 564–583.

Dandolo, L. C., & Schwabe, L. (2018). Time-dependent memory transformation along the hippocampal anterior-posterior axis. Nature Communications, 9(1), 1–11.

De Haas, N., Ottink, L., & Doeller, C. F. (2023). Integration of Euclidean and path distances in hippocampal maps. BioRxiv.

Deuker, L., Bellmund, J. L., Navarro Schröder, T., & Doeller, C. F. (2016). An event map of memory space in the hippocampus. ELife, 5, e16534.

Deuker, L., Olligs, J., Fell, J., Kranz, T. A., Mormann, F., Montag, C., … Axmacher, N. (2017). Memory consolidation by replay of stimulus-specific neural activity. In N. Axmacher & B. Rasch (Eds.), Cognitive Neuroscience of Memory Consolidation (pp. 19373–19383). Cham, Switzerland: Springer Nature.

Eichenbaum, H., Dudchenko, P., Wood, E., Shapiro, M., & Tanila, H. (1999). The Hippocampus, Memory, and Place Cells: Is It Spatial Memory or a Memory Space? Neuron, 23(2), 209–226.

Foo, P., Warren, W. H., Duchon, A., & Tarr, M. J. (2005). Do Humans Integrate Routes Into a Cognitive Map? Map-Versus Landmark-Based Navigation of Novel Shortcuts. Journal of Experimental Psychology: Learning, Memory, and Cognition, 31(2), 195–215.

Fyhn, M., Molden, S., Witter, M. P., Moser, E. I., & Moser, M.-B. (2004). Spatial representation in the entorhinal cortex. Science (New York, N.Y.), 305(5688), 1258–1264.

Gagnon, L., Kupers, R., Schneider, F. C., & Ptito, M. (2010). Tactile maze solving in congenitally blind individuals. NeuroReport, 21(15), 1.

Gagnon, L., Schneider, F. C., Siebner, H. R., Paulson, O. B., Kupers, R., & Ptito, M. (2012). Activation of the hippocampal complex during tactile maze solving in congenitally blind subjects. Neuropsychologia, 50(7), 1663–1671.

Garvert, M. M., Dolan, R. J., & Behrens, T. E. J. (2017). A map of abstract relational knowledge in the human hippocampal–entorhinal cortex. ELife, 6, 1–20.

Gaunet, F., Martinez, J. L., & ThinusBlanc, C. (1997). Early-blind subjects’ spatial representation of manipulatory space: Exploratory strategies and reaction to change. Perception, 26(3), 345–366.

Gordon, E. M., Laumann, T. O., Adeyemo, B., Huckins, J. F., Kelley, W. M., & Petersen, S. E. (2016). Generation and Evaluation of a Cortical Area Parcellation from Resting-State Correlations. Cerebral Cortex, 26(1), 288–303.

Grill-Spector, K., & Malach, R. (2004). The human visual cortex. Annual Review of Neuroscience, 27, 649–677.

Hegarty, M., Richardson, A. E., Montello, D. R., Lovelace, K., & Subbiah, I. (2002). Development of a self-report measure of environmental spatial ability. Intelligence, 30(5), 425–447.

Howard, L. R., Javadi, A. H., Yu, Y., Mill, R. D., Morrison, L. C., Knight, R., … Spiers, H. J. (2014). The hippocampus and entorhinal cortex encode the path and euclidean distances to goals during navigation. Current Biology, 24(12), 1331–1340.

Iachini, T., Ruggiero, G., & Ruotolo, F. (2014). Does blindness affect egocentric and allocentric frames of reference in small and large scale spaces? Behavioural Brain Research, 273, 73–81.

Iglói, K., Doeller, C. F., Berthoz, A., Rondi-Reig, L., & Burgess, N. (2010). Lateralized human hippocampal activity predicts navigation based on sequence or place memory. Proceedings of the National Academy of Sciences, 107(32), 14466–14471.

Iglói, K., Doeller, C. F., Paradis, A. L., Benchenane, K., Berthoz, A., Burgess, N., & Rondi-Reig, L. (2015). Interaction between hippocampus and cerebellum crus i in sequence-based but not place-based navigation. Cerebral Cortex, 25(11), 4146–4154.

Krekelberg, B., Boynton, G. M., & van Wezel, R. J. A. (2006). Adaptation: from single cells to BOLD signals. Trends in Neurosciences, 29(5), 250–256.

Kupers, R., Chebat, D. R., Madsen, K. H., Paulson, O. B., & Ptito, M. (2010). Neural correlates of virtual route recognition in congenital blindness. Proceedings of the National Academy of Sciences of the United States of America, 107(28), 12716–12721.

Lawton, C. A. (1994). Gender differences in way-finding strategies: Relationship to spatial ability and spatial anxiety. Sex Roles, 30(11–12), 765–779.

Lee, D., & Quessy, S. (2003). Activity in the supplementary motor area related to learning and performance during a sequential visuomotor task. Journal of Neurophysiology, 89(2), 1039–1056.

Loomis, J. M., Klatzky, R. L., & Giudice, N. A. (2013). Representing 3D Space in Working Memory: Spatial Images from Vision, Hearing, Touch, and Language. In S. Lacey & R. Lawson (Eds.), Multisensory Imagery: Theory & Applications (pp. 131–156). Springer.

Miao, M., Zeng, L. M., & Weber, G. (2017). Externalizing cognitive maps via map reconstruction and verbal description. Universal Access in the Information Society, 16(3), 667–680.

Morgan, L. K., MacEvoy, S. P., Aguirre, G. K., & Epstein, R. A. (2011). Distances between Real-World Locations Are Represented in the Human Hippocampus. Journal of Neuroscience, 31(4), 1238–1245.

Navarro Schröder, T., Haak, K. V., Jimenez, N. I. Z., Beckmann, C. F., & Doeller, C. F. (2015). Functional topography of the human entorhinal cortex. ELife, 4(JUNE), 1–17.

Ottink, L., Van Raalte, B., Doeller, C. F., Van der Geest, T. M., & Van Wezel, R. J. A. (2022). Cognitive map formation through tactile map navigation in visually impaired and sighted persons. Scientific Reports, 12, 11567.

Ottink, Loes, Buimer, H., van Raalte, B., Doeller, C. F., van der Geest, T. M., & van Wezel, R. J. A. (2022). Cognitive map formation supported by auditory, haptic, and multimodal information in persons with blindness. Neuroscience and Biobehavioral Reviews, 140, 104797.

Papadopoulos, K., Koustriava, E., & Kartasidou, L. (2012). Spatial Coding of Individuals With Visual Impairments. The Journal of Special Education, 46(3), 180–190.

Prestopnik, J. L., & Roskos-Ewoldsen, B. (2000). The relations among wayfinding strategy use, sense of direction, sex, familiarity, and wayfinding ability. Journal of Environmental Psychology, 20(2), 177–191.

Pruim, R. H. R., Mennes, M., Buitelaar, J. K., & Beckmann, C. F. (2015). Evaluation of ICA-AROMA and alternative strategies for motion artifact removal in resting state fMRI. NeuroImage, 112, 278–287.

Pruim, R. H. R., Mennes, M., van Rooij, D., Llera, A., Buitelaar, J. K., & Beckmann, C. F. (2015). ICA-AROMA: A robust ICA-based strategy for removing motion artifacts from fMRI data. NeuroImage, 112, 267–277.

Schinazi, V. R., Thrash, T., & Chebat, D.-R. (2016). Spatial navigation by congenitally blind individuals. Wiley Interdisciplinary Reviews: Cognitive Science, 7(1), 37–58.

Spiers, H. J., & Barry, C. (2015). Neural systems supporting navigation. Current Opinion in Behavioral Sciences, 1, 47–55.

Spiers, H. J., & Maguire, E. A. (2007). A Navigational Guidance System in the Human Brain. Hippocampus, 17, 618–626.

Stanislav, K., Alexander, V., Maria, P., Evgenia, N., & Boris, V. (2013). Anatomical Characteristics of Cingulate Cortex and Neuropsychological Memory Tests Performance. Procedia - Social and Behavioral Sciences, 86, 128–133.

Tcheang, L., Bulthoff, H. H., & Burgess, N. (2011). Visual influence on path integration in darkness indicates a multimodal representation of large-scale space. Proceedings of the National Academy of Sciences, 108(3), 1152–1157.

ThinusBlanc, C., & Gaunet, F. (1997). Representation of space in blind persons: Vision as a spatial sense? Psychological Bulletin, 121(1), 20–42.

Ungar, S. (2000). Cognitive Mapping without Visual Experience. In R. Kitchin & S. Freundschuh (Eds.), Cognitive Mapping: Past, Present and Future (pp. 221–248). Routledge, London, UK.

Viard, A., Doeller, C. F., Hartley, T., Bird, C. M., & Burgess, N. (2011). Anterior Hippocampus and Goal-Directed Spatial Decision Making. Journal of Neuroscience, 31(12), 4613–4621.

Wang, D., Qin, W., Liu, Y., Zhang, Y., Jiang, T., & Yu, C. (2014). Altered resting-state network connectivity in congenital blind. Human Brain Mapping, 35(6), 2573–2581.

Wen, Z., Zhou, F.-Q., Huang, X., Dan, D., Xie, B.-J., & Shen, Y. (2018). Neuropsychiatric Disease and Treatment Dovepress altered functional connectivity of primary visual cortex in late blindness, 3317–3327.

Wittenberg, G. F., Werhahn, K. J., Wassermann, E. M., Herscovitch, P., & Cohen, L. G. (2004). Functional connectivity between somatosensory and visual cortex in early blind humans. European Journal of Neuroscience, 20(7), 1923–1927.

Witter, M. P., Doan, T. P., Jacobsen, B., Nilssen, E. S., & Ohara, S. (2017). Architecture of the entorhinal cortex a review of entorhinal anatomy in rodents with some comparative notes. Frontiers in Systems Neuroscience, 11(June), 1–12.

Wolbers, T., Klatzky, R. L., Loomis, J. M., Wutte, M. G., & Giudice, N. A. (2011). Modality-Independent Coding of Spatial Layout in the Human Brain. Current Biology, 21(11), 984–989.

Yu, C., Liu, Y., Li, J., Zhou, Y., Wang, K., Tian, L., … Li, K. (2008). Altered functional connectivity of primary visual cortex in early blindness. Human Brain Mapping, 29(5), 533–543.

Zandbelt, B. (2017). Slice display. Figshare. 10.6084/M9.FIGSHARE.4742866

Zhang, H., Deuker, L., & Axmacher, N. (2017). Replay in Humans—First Evidence and Open Questions, 251–263.

